# The contribution of object size, manipulability, and stability on neural responses to inanimate objects

**DOI:** 10.1101/2020.11.22.393397

**Authors:** Caterina Magri, Talia Konkle, Alfonso Caramazza

**Affiliations:** Department of Psychology, Harvard University, Cambridge, MA 02138, USA; Center for Mind/Brain Sciences (CIMeC), University of Trento, 38068 Rovereto (TN), Italy

**Keywords:** object representation, occipitotemporal cortex, functional magnetic resonance imaging, real-world size

## Abstract

In human occipitotemporal cortex, brain responses to depicted inanimate objects have a large-scale organization by real-world object size. Critically, the size of objects in the world is systematically related to behaviorally-relevant properties: small objects are often grasped and manipulated (e.g., forks), while large objects tend to be less motor-relevant (e.g., tables), though this relationship does not always have to be true (e.g., picture frames and wheelbarrows). To determine how these two dimensions interact, we measured brain activity with functional magnetic resonance imaging while participants viewed a stimulus set of small and large objects with either low or high motor-relevance. The results revealed that the size organization was evident for objects with both low and high motor-relevance; further, a motor-relevance map was also evident across both large and small objects. Targeted contrasts revealed that typical combinations (small motor-relevant vs. large non-motor-relevant) yielded more robust topographies than the atypical covariance contrast (small non-motor-relevant vs. large motor-relevant). In subsequent exploratory analyses, a factor analysis revealed that the construct of motor-relevance was better explained by two underlying factors: one more related to manipulability, and the other to whether an object moves or is stable. The factor related to manipulability better explained responses in lateral small-object preferring regions, while the factor related to object stability (lack of movement) better explained responses in ventromedial large-object preferring regions. Taken together, these results reveal that the structure of neural responses to objects of different sizes further reflect behavior-relevant properties of manipulability and stability, and contribute to a deeper understanding of some of the factors that help the large-scale organization of object representation in high-level visual cortex.

**Highlights:**

- Examined the relationship between real-world size and motor-relevant properties in the structure of responses to inanimate objects.
- Large scale topography was more robust for contrast that followed natural covariance of small motor-relevant vs. large non-motor-relevant, over contrast that went against natural covariance.
- Factor analysis revealed that manipulability and stability were, respectively, better explanatory predictors of responses in small- and large-object regions.

## 1. Introduction

To recognize objects in the real world, large portions of the brain are utilized, including the ventral visual stream (Goodale and Milner, 1992; Ishai et al., 1999). Within this cortex there is a highly consistent organization of neural response tuning, with a tripartite large-scale topography distinguishing responses to animate entities, large inanimate objects, and small inanimate objects (Chao et al., 1999; Konkle and Caramazza, 2013; Proklova et al., 2016; Julian et al., 2017; Grill-Spector and Weiner, 2014). Some proposals have raised the possibility that the large-scale divisions in this cortex arise to support different behavioral needs (e.g., manipulating objects vs navigating in environments), linked to distinct underlying long-range brain networks (e.g., Mahon and Caramazza, 2011; Konkle and Caramazza, 2017). Thus, in the present work we examined how both the size of an object and its motor-relevance can explain brain responses to object categories.

Intuitively, the relationship between an object’s size and the degree of interaction it affords in the world is evident: small objects tend to be graspable and easy to hold up, while larger objects tend to be used as support or for navigation. However, other relationships are possible, including not usually manipulated small objects like picture frames, and interactive large objects like pianos. Here, we operationalize the dimension dealing with object interaction as *motor-relevance*. While similar conceptually to the property of manipulability that has been investigated in the past (Kellenbach et al., 2003; Boronat et al., 2003; Mahon et al., 2007; Campanella et al., 2010; Kalénine and Buxbaum, 2016), motor-relevance spans a wider definition in that it includes actions performed not only with the hands, but with other parts of the body as well. With this broader conceptualization, size can more easily dissociate from degree of interaction: large objects like swing sets and wheelbarrows tend to involve movements beyond hand-performed actions, while for small objects such as picture frames and smoke alarms there is no clear motor interaction associated.

To date, there is substantial evidence that motor-relevance is an important construct for visual object responses, particularly along the lateral aspect of the occipitotemporal cortex, a region known to show a preference for small inanimate over large inanimate objects (Konkle and Oliva, 2012; Konkle and Caramazza, 2013). For example, in this territory, there is a region with preference for tools (Chao et al., 1999; Bracci et al., 2012; Gallivan et al., 2013; Chen et al., 2016) which is closely overlapping with effector-specific regions (Bracci et al., 2012), and persists in the congenitally blind (Peelen et al., 2013; Wang et al., 2015). Further, this cortex shows connectivity with frontoparietal networks supporting actions (Simmons and Martin, 2012; Bi et al., 2015; Konkle and Caramazza, 2017), and is causally involved in tool-action discrimination judgments (Perini et al., 2014). While past studies have typically focused on objects that require hand-performed actions, the present study allows us to test whether the inclusion of non-hand-performed actions through the motor-relevance dimension would further explain neural responses and extend them to large objects.

In the ventromedial temporal cortex, another tool-preferring region has been observed by some studies in the medial fusiform gyrus when contrasting tools to animals (Chao et al., 1999; Whatmough et al., 2002; Mahon et al., 2007; Garcea and Mahon 2014). However, the property of motor-relevance in this region is less clear: for example, responses to manipulable tools are not higher than to other inanimate objects (Mahon et al., 2007; Chen et al., 2018), though there is still some evidence for sensitivity to tools in neural adaptation signals (Mahon et al., 2007). Further, research focused on scene understanding has characterized these ventral-medial object responses along other kinds of object properties that are less related to motor-relevance. For example, responses in ventromedial scene-related regions are best predicted by objects invoking a local space that tend to stay fixed in the world, and are useful as a landmark (Troiani et al., 2014, Mullally and Maguire, 2011; Julian et al., 2016; Auger et al., 2013; see also Epstein, 2014). These same areas are also known to show a preference for large over small inanimate objects (Konkle and Oliva, 2012; Konkle and Caramazza, 2013) and, interestingly, also to the names of such objects in both blind and sighted individuals (He et al., 2013).

Given these past observations, in the present study we aimed to clarify how brain responses to different objects in occipitotemporal cortex reflect motor-relevance and object size properties. To do so, we designed our experiment to enable two different levels of granularity. First, we compared the broad relationship among object size and motor-relevance properties along a 2×2 design (large vs small, motor-relevant vs non-motor relevant), finding effects for both variables independently. Second, we examined brain responses at the level of object categories (72 categories total), where a factor analysis revealed that three dimensions that respectively relate to object size, manipulability, and stability interact to account for the structure of brain responses in different regions. Finally, we consider the relative merits of visual feature-versus domain-based properties (e.g., manipulability, navigation relevance) as driving principles in the organization of high-level visual cortex.

## 2. METHODS

### 2.1 Participants

For the fMRI experiment, 21 healthy adults with normal or corrected-to-normal vision participated in a 2-hour fMRI session (age, 18-30 years; 11 females). Three were later removed for excessive head movement, and the remaining 18 subjects were analyzed. Each subject gave informed consent according to procedures approved by the Institutional Review Board at the University of Trento.

For the online behavioral experiments, ratings were collected from 644 total participants (464 subjects for the stimulus set development and validation over the dimensions of motor-relevance [n=161], size [n=168] and recognition [n=135]; and 36 subjects for each of the dimensions of manipulability, spatial definition, position variability, motion-based identity and interaction envelope).

### 2.2 Stimuli

The stimulus set consisted of images from 72 object categories, where each object category belonged to one of four conditions (small motor-relevant, small non-motor-relevant, large motor-relevant and large non-motor-relevant) with 18 object categories per condition. Each object category included images of six exemplars of that category, for a total of 432 pictures in the stimulus set (Figure 1a). Each image depicted an isolated object on a white background, and the image was sized to 512 x 512 pixels with the object’s longest axis reaching the border of the image.

**Figure 1.**
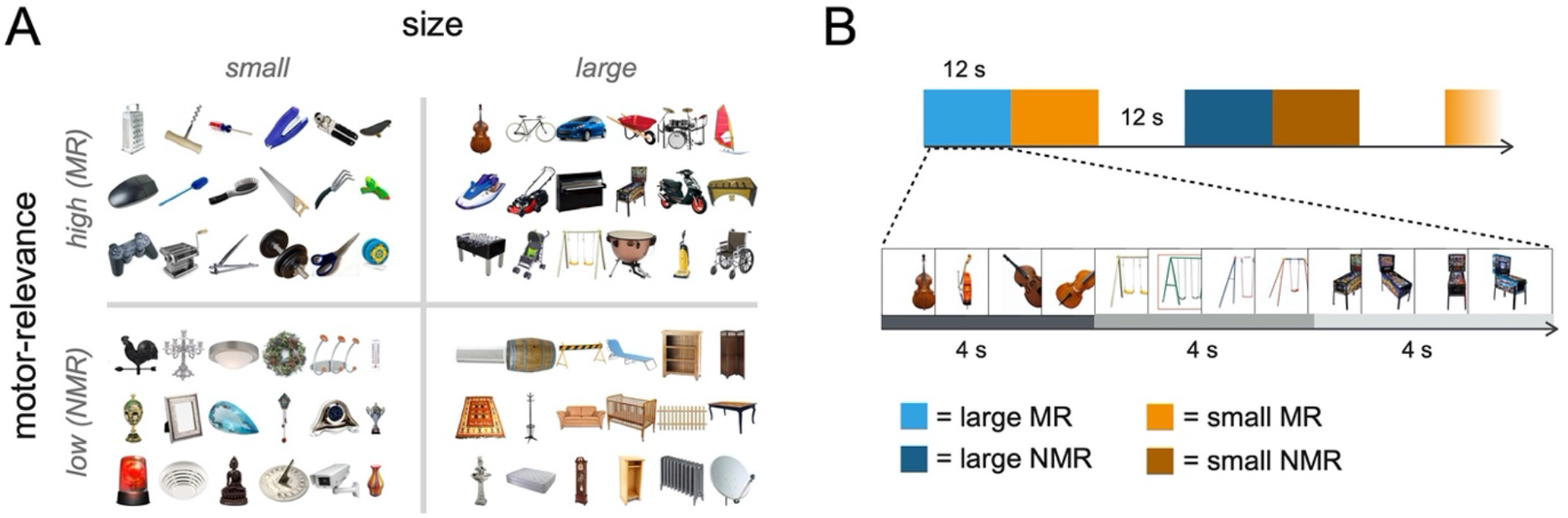
a) Stimulus set: 72 inanimate object categories divided into four conditions (small motor-relevant, small non-motor-relevant, large motor-relevant, large non-motor-relevant). Each object category included six individual items, for a total of 432 individual images. b) fMRI task: subjects were presented with four consecutive pictures of the same object category in a 4s object category mini-block. Three 4s category-level mini-blocks (e.g. cello mini-block, swing set mini-block, pinball machine mini-block) were presented sequentially in a 12s condition-level main block (e.g. large motor-relevant objects block). The task for participants was to detect when a red frame surrounded one of the objects, which happened once per 12s block.

#### 2.2.1 Stimuli validation

To develop and validate the stimulus set, we performed three separate behavioral studies using Amazon Mechanical Turk on an initial larger set of 20 object categories per condition with 7 unique exemplars for each object category (560 total images). Our aim was to construct a stimulus set in which items were recognizable, and where the size and motor-relevance dimensions were balanced.

a. *Recognition ratings*. Observers were presented with an item and reported whether they were familiar with this object (yes/no), and how easily they recognized the object on a scale from 1 (very hard to recognize) to 5 (very easy to recognize).
b. *Motor-relevance ratings*. Participants were presented with an item and rated the degree to which the object made them think of moving their hands or other parts of their body, using a scale from 1 (not at all) to 7 (very strongly).
c. *Size Ratings*. Participants were presented with an item and judged the size of an object as they would encounter it in everyday life, on a scale from 1 to 7. To provide a reference for the scale, pictures of a wristwatch (small size), a hiking backpack (medium size) and a bed (large size) were provided beside the values of 1, 4 and 7 on the scale.

For all three experiments, each participant made judgments for 20 items (5 items from each of the four conditions). Only one exemplar for an object category was presented to each participant. Ratings were obtained for all 560 objects combining data from all participants, so that each image was rated by 6 observers along the measures of size and motor-relevance and by five observers along the measure of recognition.

In order to eliminate those items that were least recognizable, we dropped one item from each object category based on the familiarity score in the recognition experiment. We next eliminated two object categories that scored low in familiarity in order to balance the ratings from the other two experiments (Motor-relevance and Size experiments). This resulted in a new stimulus set with 18 objects per condition (72 total) and 6 items per object. All results and analysis that follow refer to this new stimulus set.

To ensure these factors were balanced, we conducted several statistical tests. As expected, motor-relevant objects had higher motor-relevance scores than non-motor-relevant objects (t_35_=32.93, p<0.001), and large objects had higher size ratings than small objects (t_35_=19.82, p<0.001). Further, large and small objects did not differ in their average motor-relevance scores (t_35_=1.63, p=0.11). The same was true when looking separately at large motor-relevant and small motor-relevant objects (t_17_=1.05, p=0.31), and small non-motor-relevant and large non-motor-relevant object categories (t_35_=1.35, p=0.2). Similarly, motor-relevant and non-motor-relevant objects did not differ in their average size score (t_35_=-0.72, p=0.5) and neither did large motor-relevant and large non-motor-relevant relevant objects (t_17_=0.96, p=0.35), nor small motor-relevant and small non-motor-relevant objects (t_17_=-1.76, p=0.1).

### 2.3 Image Acquisition

Imaging data were acquired using a BioSpin MedSpec 4T scanner (Bruker) with an eight-channel coil. Functional data were obtained with an echo-planar 2D imaging sequence (repetition time TR: 2000ms; echo time TE: 33ms; flip angle: 73º; slice thickness: 3 mm; gap: 0.99 mm, with 3×3 in-plane resolution and 34 slices). Volumes were acquired in ascending interleaved order of slice acquisition.

### 2.4 Experimental Design and Statistical Analysis

#### 2.4.1 Design and procedure

During the fMRI experiment subjects viewed images of the small motor-relevant, small non-motor-relevant, large motor-relevant, large non-motor-relevant objects in a classic blocked design. In each run, stimuli from each of the 4 conditions were presented 6 times in 12s stimulus blocks, with 6 rest blocks of 12s interspersed. Images were each presented for 800ms with a 200ms blank between each stimulus. Fixation blocks also appeared at the beginning and end of each run, for 4 and 12 seconds respectively. Participants were instructed to maintain fixation and to press a button when a red frame surrounded one of the objects, which happened once per block. The full experiment consisted of six runs each lasting 6 minutes and 12 seconds.

Additionally, we added sub-structure in each stimulus block, to support more exploratory analyses about the relationship between all 72 object categories (see Figure 1b). Specifically, each stimulus block was comprised of 4s mini-blocks. In each mini-block, three different exemplars from the same category were shown. For example, in a 12s block of large motor-relevant objects, a participant would see 4 different cellos, followed by 4 different swings, followed by 4 different pinball machines. Within each run all 72 object categories were presented exactly one time. The specific 4 exemplars that were displayed in each mini-block were randomly chosen without repetition from the set of 6 possible exemplars from that object category.

Each image was presented in isolation at a ∼8 x ∼8º visual angle. Stimulus presentation was performed using the Psychophysics Toolbox package (Brainard, 1997) in MATLAB.

#### 2.4.2 Functional Localizer

Three independent functional localizer runs were performed to identify category-selective regions. These runs consisted of a blocked design, in which each block included one of the following stimuli: 1) objects 2) scrambled objects 3) scenes 4) bodies 5) hands. Each run lasted 6 minutes and 36 seconds, with 50 6-second stimulus blocks (10 per condition) and ten 8-second rest blocks interspersed. In each stimulus-block, six different stimuli from the same condition were presented, where each image was shown for 800ms followed by 200ms of fixation. Additionally, there was an orthogonal motion manipulation: for each of the 5 stimulus categories, in half of the blocks the object was presented at the center of the screen, and in the other half the object moved from the center out in one of 8 randomly directions randomly. Observers were instructed to maintain fixation throughout the experiment and press the button when an exact image repeated back-to-back.

Two of the 21 subjects performed 8 experimental runs (instead of 6) and only 2 Localizer runs (instead of 3). Data from all experimental runs for these subjects were used for data analysis. Due to a technical error, localizer runs for one subject did not include scrambled objects.

For the purpose of the current project, we were only interested in identifying category-selective regions Lateral Occipital Complex (LOC; Objects > Scramble; Grill-Spector, 2003), parahippocampal place area (PPA; Scenes > Objects; Epstein and Kanwisher, 1998) and Occipital Place Area (OPA; Scenes > Objects; Dilks et al., 2013). We were not able to localize area LOC in the subject for whom the scrambled objects condition was missing. To compare the coordinates of our contrasts of interest with the location of well-known category-selective regions, we also localized extrastriate body area (EBA; Downing et al., 2001) and a nearby hand-selective area (Bracci et al., 2012).

#### 2.4.3 fMRI Data Analysis

Functional neuroimaging data were analyzed using SPM12 (Ashburner et al., 2014), MARSBAR (Brett et al., 2002) and bspmview (https://www.bobspunt.com/software/bspmview/) on MATLAB (Versions 2014b and 2016a,b). Barplots and scatterplots were displayed using ggplot2 (Wickham, 2009) in RStudio.

The raw functional images were submitted to preprocessing, where the first 4 volumes were discarded from each run, slice scan-time correction was performed, followed by 3D motion correction, normalization and spatial smoothing (8 mm FWHM kernel). Data were modeled using standard general linear models (GLM). The first GLM included regressors for each of the four main conditions, with run regressors and motion correction parameters included as nuisance factors. The second GLM modeled each object category as a separate condition for a total of 72 regressors, with run regressors and motion correction parameters included as nuisance factors.

#### 2.4.4 Regions-of-interest selection

We created our ROIs utilizing a separate set of runs from the ones we used for data-analysis. Spherical ROIs were defined around the individual peaks of activation for the whole-brain size contrast Small > Large (collapsing over motor-relevance) from two experimental runs. Univariate responses were extracted from the remaining experimental runs for size and motor-relevance conditions. These ROIs were defined with a 12mm sphere (257 voxels) centered around the peak positive and negative voxels in individual subjects, and then intersected with individual whole-brain gray matter masks. The selected peaks are referred to following the conventions of Konkle and Oliva (2010) and included small-object preferring bilateral inferior temporal gyrus (Small-ITG, left hemisphere [LH] n=15, right hemisphere [RH] n=17), and large-object preferring bilateral PHC (Large-PHC, LH n=16, RH n=15). After intersecting the 12mm ROI sphere with each subjects’ gray-matter mask, the ROIs slightly varied in size across subjects. After intersection, the average size was 244 voxels for Small-ITG and 257 voxels for Large-PHC.

We further identify large-object preferring TOS (Large-TOS) dorsally based on the size contrast, and a series of category-selective regions using the Functional Localizer runs. The regions identified with the Localizer were object-selective lateral occipitotemporal cortex (LOC), scene-selective parahippocampal place area (PPA), and scene-selective occipital-place area (OPA). Analysis and results for Large-TOS and the category-selective ROIs are reported in the Supplementary Material.

#### 2.4.5 Voxel mask

We produced a mask of the most reliable object-selective voxels in occipital, temporal and parietal regions using the reliability-based voxel selection method for condition-rich designs (Tarhan and Konkle, 2020). To compute the reliability maps, average beta values for odd and even runs were separated in two groups and correlated for each voxel that was included within the gray matter. To establish a reliability threshold, we then correlated the patterns for the same object category at different thresholding levels and picked the threshold at which the average correlation across conditions plateaued, which in this case was r = 0.25. We then produced a mask from the reliability map at this value.

#### 2.4.6 Whole-brain conjunction analysis

To explore the size organization across motor-relevance, we performed a random-effects conjunction analysis at the individual level between small motor-relevant > large motor-relevant and small non-motor-relevant > large non-motor-relevant. For each individual, at each voxel, t-values for the two contrasts were compared. The β-weight corresponding to the contrast with t-value closest to zero was then assigned to the given voxel. This resulted in eighteen conjunction maps (one for each participant) which were then submitted to a one-sample t-test, and the resulting t-values were assigned to the corresponding voxels on a new map. The voxels with highest value in these new t-maps are those that are strongly activated or deactivated by both contrasts in most subjects. The resulting maps were thresholded with q(FDR)<0.05. Similarly, to explore the motor-relevance organization across size, we performed the same analysis but considering small motor-relevant > small non-motor-relevant and large motor-relevant > large non-motor-relevant contrasts.

#### 2.4.7 Typical-Atypical contrast analysis

To further explore the relationship between size and motor-relevance, we compared maps between two contrasts: a *typical contrast* [small motor-relevant > large non-motor-relevant], which follows the real-world covariation of small objects being more motor-relevant than large objects; and an *atypical contrast* [small non-motor-relevant > large motor-relevant]. Individual maps for each contrast were submitted to a second-level analysis to obtain a group map, which was FDR-thresholded at q<0.05.

To assess the reliability of the two contrasts, we further produced individual maps for each contrast separately for odd and even runs and correlated them. We then submitted individual correlations for the two separate contrasts to a t-test to assess whether there was a significant different in reliability between the maps produced by the two contrasts.

### 2.5 Exploratory Analysis

#### 2.5.1 Object Property Ratings

Ratings for the 72 objects in our dataset were collected on a series of other dimensions to identify the ones that were best associated with the response profiles of our ROIs. All studies were conducted on Amazon Mechanical Turk with 36 judgments per object category. Each of the participants (n=36) rated one exemplar of each of the 72 object categories, with six participants rating the same exemplars within each category. Participants were not presented with two exemplars from the same category. The following object properties were rated:

1. *Manipulability*. Manipulability tracks the degree to which hand-interactions with objects are prominent. Participants were presented with an item and rated the degree to which the object made them think of moving their hands, using a scale from 1 (not at all) to 7 (very strongly). The question asked to participants was very similar to the one posed to collect the motor-relevance dimension, but here we focus on actions performed with the hands rather than with other parts of the body.
2. *Motion-based Identity*. The degree to which an object’s parts are expected to move and change position might be a relevant property for representing objects. Participants were asked to rate on a scale from 1 to 7 the degree to which motion is important in determining the identity of the object.
3. *Spatial Definition*. This property is defined as “the degree to which objects evoke a sense of surrounding space” (Mullally and Maguire, 2011). Participants were asked to rate on a scale from 1 to 7 the degree to which a background is evoked when looking at a specific object.
4. *Position Variability*. Objects can be more or less likely to change location in the environment (i.e., a car is more likely to change location than a swing). Participants were asked to rate on a scale from 1 to 7 the degree to which an object is likely to change position in everyday life.
5. *Interaction Envelop*e. “Interaction envelope” was operationalized as a measure of the amount of space needed to interact with the object (e.g., a hamburger needs to be manipulated with two hands that cover its whole surface, thus it has a larger interaction envelope than a coffee mug which just needs one hand on its handle to be manipulated), following Bainbridge and Oliva (2015). Participants had to judge on a scale from 0 to 2 how many hands are often used when interacting with an object.

#### 2.5.2 Pairwise correlations

We compared the similarity across our seven dimensions (size, motor-relevance and the additional ones collected post-hoc), and the reliability of each of the rated dimensions, in the following way. First, we divided participants into two groups and computed the mean rating for each group and each object category separately for each dimension, resulting in two group vectors of 72 mean ratings for each dimension. To compute the reliability of a dimension, we correlated the two group vectors with each other. To compute the correlation between two different dimensions, we correlated each dimension’s group vectors with the other dimension’s group vectors, and averaged across the four combinations.

#### 2.5.3 Factor Analysis

To understand how our critical dimensions of size and motor-relevance related to the variety of stimulus properties from the subsequent stimulus norming studies, we conducted a maximum-likelihood factor analysis on all seven dimensions (size, motor-relevance, manipulability, motion-based identity, spatial definition, position variability and interaction envelope) with a “varimax” rotation of the coefficients. This factor analysis was implemented with the *factanal* function in R. To determine the number of meaningful factors to extract from our dimensions, we performed a “parallel” analysis (Horn, 1965). This procedure involves generating random data and submitting them to the same factor analysis, iterated 5000 times. The average of the eigenvalues resulting from the parallel random-data factor analyses is then compared to the factors from the observed data: if the average eigenvalues from the parallel factor is smaller than the one from the data factor, then the data factor is kept. This procedure was implemented with the *fa*.*parallel* function in R. The three resulting factors can be summarized as relating to real-world size and context (f_size_), manipulability (f_manip_) and degree of movement (f_stability_). For the following ROI and searchlight analyses, for ease of interpretation, we flipped the direction of f_stability_ so as to positively correlate with degree of stability.

#### 2.4.6 ROI analysis with Factors as predictors

Following the factor analysis on the behavioral ratings of the objects, we performed a linear model analysis to predict the size ROIs responses from the resulting factor scores and their interactions. To estimate the response profile of each ROI, β-weights for each of the 72 object categories were extracted from the GLMs of the main experiment runs. Note that this analysis was performed on runs separate from the ones used to produce the ROIs. These betas were averaged across voxels, resulting in a vector of 72 values reflecting the ROIs overall response activation to each category. The ROIs activation profiles were computed for each subject using subject-specific ROIs and GLMs.

We next sought to explain these ROI activation profiles with the factors from the factor analysis above by employing a linear modeling approach. All linear models to predict an ROI activation were implemented with the function *lm* in R. We first looked for hemispheric interactions by submitting our size ROIs to a model with the three factors, hemisphere and their interaction as coefficients. No hemispheric interaction significantly explained either Small-ITG or Large-PHC’s activation; hemisphere was thus removed from the models.

Next, we examined which of three nested models best explained activation in our size ROIs by comparing their resulting adjusted R square (adjR^2^). The three models were: one with the three factors as predictors, with no interaction terms [f_size_ + f_manip_ + f_stability_]; a second model that also included the two-way interaction terms [f_size_*f_manip_ + f_size_*f_stability_ + f_manip_ *f_stability_]; and a third model that also included the three-way interaction term [f_size_*f_manip_*f_stability_]. The model with the highest adjR^2^ was considered the one most fitting the region and the significant coefficients were explored. If two models had comparable adjR^2^ (rounded at the second decimal), the simpler model was picked. The same analysis was also performed on Large-TOS and in category-selective regions defined from the Functional Localizer (LOC-sphere, PPA-sphere and OPA-sphere; see Supplementary Material).

When comparing the nested models for the highest adj R^2^, we found that both Small-ITG and Large-PHC were best explained by a model with all three factors and their two-way interactions (Small-ITG: adj. R^2^ = 0.33; F_6,65_ = 6.90, p < 0.001; Large-PHC: adj. R^2^ = 0.27; F_6,65_ = 5.36, p < 0.001), as compared to the model with only main effects (Small-ITG: adj. R^2^ = 0.23; Large-PHC: adj. R^2^ = 0.21) and the model with main effects, two- and three-way interactions (Small-ITG: adj. R^2^ = 0.32; Large-PHC: adj. R^2^ = 0.26).

To evaluate the quality of the fit of the different coefficients, we looked at the resulting associated t- and p-values. T-values were computed by dividing coefficients by their standard errors, while p-values tested the significance against the hypothesis of obtaining given t-values if the coefficients were not actually contributing in explaining the dependent variable.

#### 2.5.4 Searchlight Regression Analysis

Following our findings in the spherical ROIs, we next explored which factors would best fit voxels throughout object-responsive cortex.

The searchlight analysis was performed in the following way: within the voxel mask, a 6mm sphere was drawn around a voxel, and all voxels falling within that sphere were considered part of the neighborhood for that voxel. This procedure was performed iteratively so that each voxel within the object-selective mask was at the center of the sphere. In each searchlight ROI, voxel activation for all 72 object categories was averaged across all voxels within the sphere, and the resulting= vector used as the dependent variable for two models.

For each search sphere, we fit two models. The first model related size and manipulability, with f_size_, f_manip_, and their interaction [f_size_ + f_manip_ + f_size_ * f_manip_], which we refer to as *manip-model*. the second model related size and stability, with f_size_, f_stability_ and their interaction [f_size_ + f_stability_ + f_size_ * f_stability_], which we refer to as *stability-model*. For each given searchlight, adjusted R^2^ values for these two models were extracted and compared, and the model with the highest positive value was selected and assigned to that searchlight’s central voxel.

To visualize which models are fitting best and where, we looked separately at small-preferring and large-object preferring voxels. Small- and large-object preferring voxels were identified based on whether the coefficient for the main effect of f_size_ was positive (preference for small) or negative (preference for large). We then visualized the coefficient maps for the interaction terms f_size_*f_manip_ and f_size_*fs_tability_ separately for large- and small-preferring voxels, comparing them with a map of the F-values for the winning model associated with each voxel.

## 3. RESULTS

### 3.1 Whole-brain conjunction analysis

We first examined whether the large-scale neural organization by real-world size of inanimate objects is preserved when disrupting the natural covariance between size and motor-relevance. To do so, we conducted a conjunction analysis to isolate regions which show a large vs small object difference that holds when comparing motor-relevant as well as non-motor-relevant objects (see Methods).

When looking at the real-world size organization, we found regions that showed a preference for small over large objects, and for large over small objects, regardless of the motor-relevance preference. The whole brain maps can be visualized in Figure 2a, with the group MNI coordinates at the bottom of the figure. With respect to small objects, we observed activation bilaterally in ITG and in the left hemisphere and along the lateral aspect of the fusiform gyrus. In the left hemisphere, the region for small objects expanded posteriorly to the lateral occipital cortex. With respect to large objects, along the ventral surface of the occipitotemporal cortex, we observed bilateral regions corresponding to the parahippocampal cortex. A preference for large objects was also present more dorsally in the lateral surface in the occipital place area (OPA) in individual participants, but this region did not survive the group-level conjunction test at the FDR threshold depicted in Figure 2a. Overall, this group-level conjunction analysis reveals that, even when taking into account motor-relevance, the real-world size of objects drives large-scale differential responses across the ventral stream.

**Figure 2.**
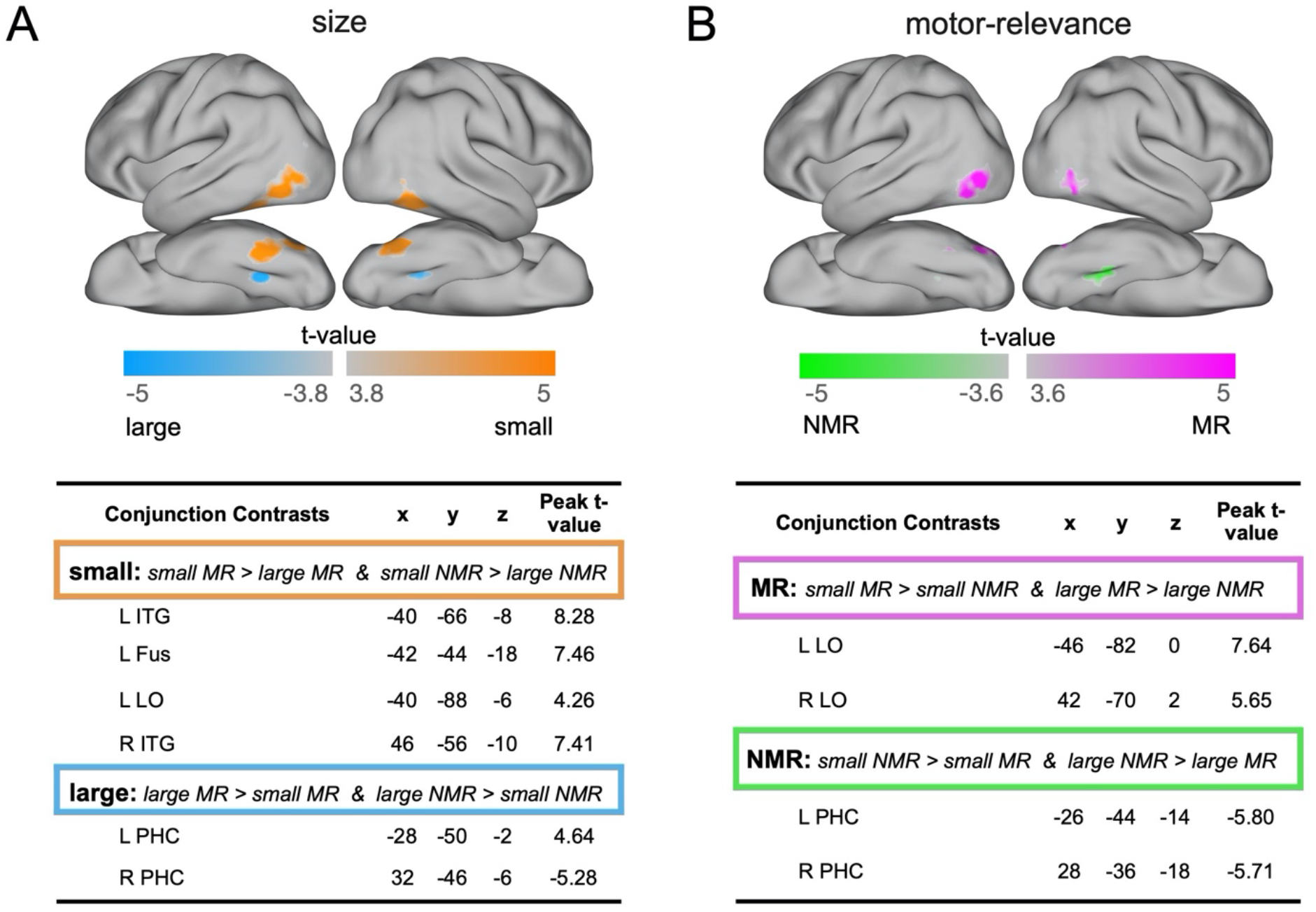
Whole Brain Random-Effects Conjunction Analysis. a) Size conjunction contrasts, where both [small motor-relevant > large motor-relevant] and [small non-motor-relevant > large non-motor-relevant] relationships hold. a) Motor-relevance conjunction contrasts, where both [small motor-relevant > small non-motor-relevant] and [large motor-relevant > large non-motor relevant] relationships hold. Contrasts were thresholded at q(FDR)<0.05.

Next, we examined whether also motor-relevance, when disrupting its natural covariance with size, would elicit a large-scale organization across this cortex. Another whole-brain conjunction analysis was conducted, comparing motor-relevant objects vs non-motor-relevant objects, requiring this relationship to hold for both large object and small objects independently. The results are mapped in Figure 2b, with the group MNI coordinates at the bottom of the figure. Along the lateral surface, we observed regions with a stronger response to motor-relevant objects, with peaks in LO bilaterally. Along the medial aspect of the ventral surface, we found stronger responses to non-motor-relevant objects primarily in the right hemisphere, with the peak activation located slightly anterior to the large-object preferring region. This analysis reveals that the dimension of motor-relevance also elicits stable response differences in parts of object-responsive cortex.

Notably, these conjunction maps yielded relatively similar organizations. Specifically, lateral regions prefer both small and motor-relevant objects; while ventromedial regions prefer both large and non-motor-relevant objects. These findings indicate that neither size alone nor motor-relevance alone are sufficient to explain the large-scale organization of responses to objects along occipitotemporal cortex. Further, these results highlight that there is a clear association between the two dimensions and how they map across the cortex.

### 3.2 Typical-atypical Contrasts

To further explore the relationship between size and motor-relevance, we compared maps between two contrasts: a *typical contrast* (small motor-relevant > large non-motor-relevant), which follows the real-world covariation of small objects being more motor-relevant than large objects; and an *atypical contrast* (small non-motor-relevant > large motor-relevant; Figure 3a). These maps are shown in Figure 3.

**Figure 3.**
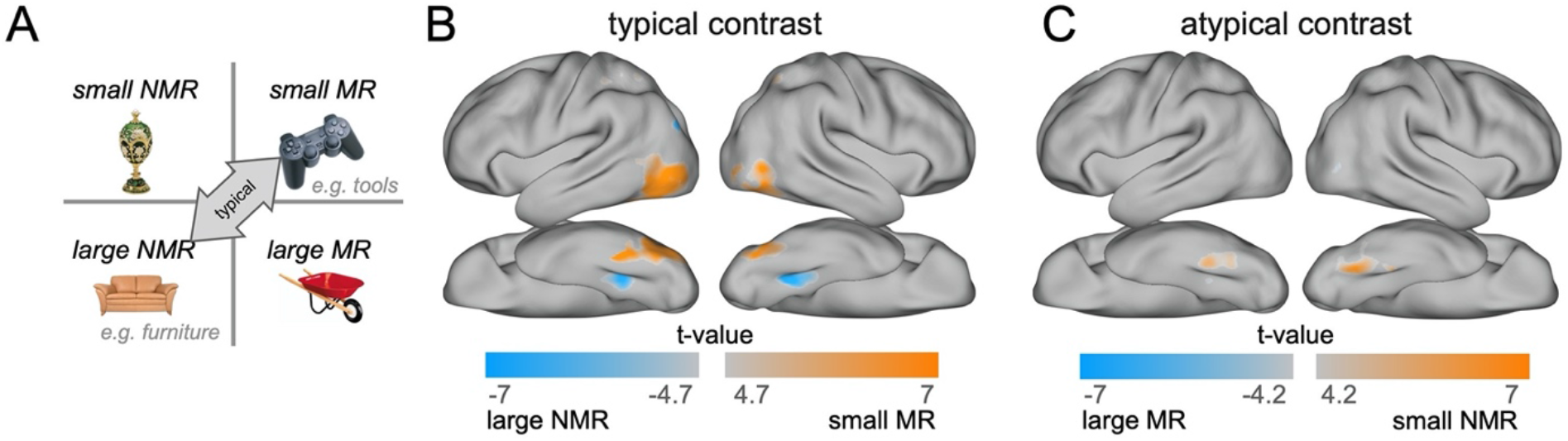
Typical/Atypical contrast Analysis. a) simplified representation of the size x motor-relevance design. Arrow points to the typical contrast conditions, representing the covariation commonly observed in the environment. b) Typical contrast: small motor-relevant > large non-motor-relevant. c) Atypical contrast: small non-motor-relevant > large motor-relevant. Contrasts were thresholded at q(FDR)<0.05.

When looking at surface maps produced by the typical contrast, we find extensive activations along the lateral and ventral surfaces (Figure 3b). In contrast, the surface maps produced by the atypical contrast (Figure 3c) are less extensive and also less reliable in a split-half analysis (typical split-half map correlations: M=0.61, SD=0.18; atypical: M=0.43, SD=0.24; t_17_=2.70, p<0.05).

These topographic observations mirror the ecological covariation of these factors (Figure 3a): while it is theoretically possible to artificially dissociate size and motor-relevance as we have done in this stimulus set, in every day experience small objects tend to be more motor-relevant and large objects less so. And we found this covariation to also be evident in the neural data (see Figure 3b,c). Taken together, these data show the importance of the interaction between size and motor-relevance in driving more reliable and robust neural responses, thus revealing the importance of the natural covariation between these two properties in the visual system.

### 3.3 Exploratory Analyses

#### 3.3.1 Factor Analysis

We next explored the possibility that the relationship between size and motor-relevance may be more multifaceted than what can be gathered from a simple 2 x 2 design. Our goal for this analysis was to understand and explore how other, related properties that have been proposed in the past relate to our findings and predict neural responses at the category level. To do so, we collected ratings along five additional dimensions of our 72 object categories and submitted them to a factor analysis together with size and motor-relevance.

First, manipulability ratings were obtained, which focused on hand movements specifically, in order to understand the extent to which our motor-relevance dimension correlated with this previously studied dimension (Mahon et al., 2007). We also examined a number of dimensions that have been previously proposed as significant modulators of tool regions or scene regions, namely interaction envelope (Bainbridge and Oliva, 2015), spatial definition (Mullally and Maguire, 2011), and motion-based identity (Beauchamp, 2005). Finally, based on our observation regarding low responses to vehicles in Large-PHC, we obtained “position variability” ratings, capturing how often an object changes its position in the environment. Note that these properties were defined after having seen the data, and thus any role they play in explaining neural data requires independent follow-up studies to confirm.

The relationship among these ratings and our original size and motor-relevant dimensions are detailed in Figure 4. All ratings had a relatively high reliability (Figure 4a, shaded diagonal numbers), with a range of similarity relationship between any pair of dimensions (Figure 6b). There are a couple observations to note from inspecting these pairwise correlations and scatter plots. First, by design, size and motor-relevance were completely uncorrelated in this stimulus set. Second, the correlation between motor-relevance and manipulability was at ceiling, virtually identifying the same property. As mentioned in the introduction, most motor-relevant objects involved hands for their interaction; including whole-body actions in the motor-relevance score had a minimal or non-existent effect (the only exception being the skateboard which was high in motor-relevance but low on manipulability).

**Figure 4.**
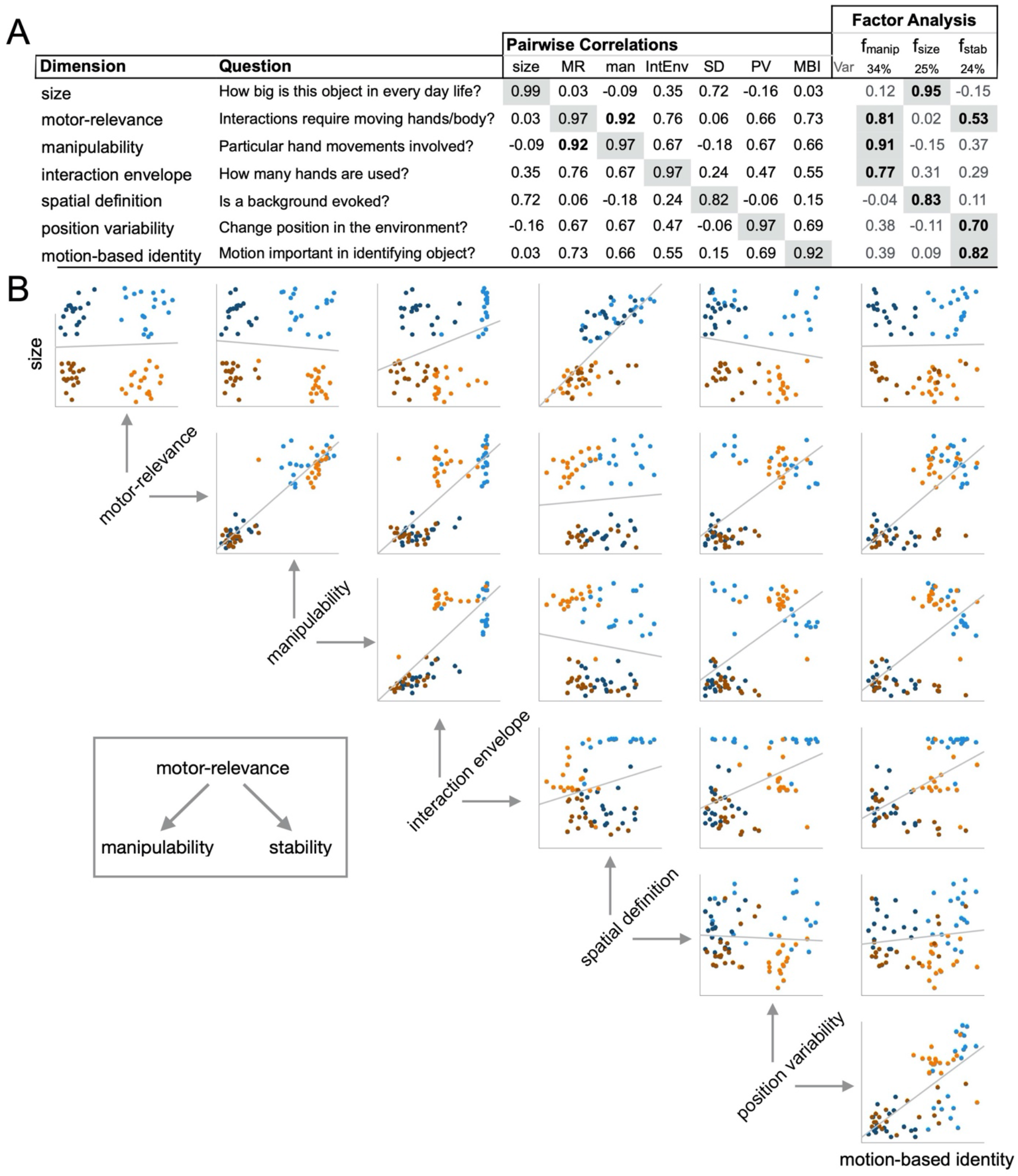
Object dimension similarity and Factor Analysis. a) For each rated object property, the subplot shows the dimension label, the question asked to elicit ratings on the dimension, the correlation to the other dimensions, where the rating’s reliability is shaded, and the dimension’s loading in a factor analysis. b) Scatter plots are shown for all pairs of 7 dimensions, where each dot reflects an object category and is color-coded based on its size and motor-relevance rating (light orange – small motor-relevant; dark orange: small non-motor relevant; light blue: large motor-relevant; dark blue: large non-motor-relevant).

Next, we extracted the latent dimensions within this dataset using factor analysis (see Methods). A parallel analysis procedure determined that three factors should be retained, which were sufficient to explain more than 80% of all variance among the ratings. The property loadings for each factor are shown to the right in Figure 4a.

The first factor was loaded mostly by manipulability, motor-relevance and interaction envelope—it seemed thus to be most related to *actions and hand-object interactions*. The second factor was loaded by size and spatial definition, and was thus most related to the *physical presence* of the object. Finally, motion-based identity, position variability and motor-relevance loaded strongly on the third factor, which seemed thus to capture the *mobility/movability of the object*.

Considering these factors within the context of our main design, our motor-relevance dimension was effectively split into two factors, one related to hand-object interactions (f_manip_) and one related to an object’s mobility/movability (f_stability_), and both of these were separate from the more physical properties of size and, to a lesser degree, spatial definition (f_size_). Thus, while our stimulus set was collected as varying along only two primary factors (size and motor-relevance), the factor analysis indicated that this stimulus set is better characterized along three factors (size, manipulability and stability).

We next explored how well the three factors extracted from our ratings can explain the neural response profiles, both in a targeted ROI analysis and in a broader searchlight analysis. Note that these analyses were unplanned and should be considered post-hoc and exploratory in nature.

#### 3.3.2 ROI Analysis

To explore how well the three factors could explain fMRI responses, we employed a linear modeling analysis explaining the activation profile in each ROI, with the three factors obtained from the factor analysis and their interactions as regressors (see Methods). The goal was to see which weighted combination of factors best predicted a region’s response variation to the 72 objects. This finer-grained analysis looking at responses to all 72 object categories was possible due to our nested fMRI protocol design (see Methods), in which the neural responses could be modeled at the category-level for each of the 72 categories independently (see Figure 1). Note that these three factors were determined only from the behavioral ratings of our stimulus set. Further, for interpretive clarity, we reversed the sign of f_stability_ so that high values indicate high “fixedness” (rather than high mobility.) This transformation does not affect the significance of the statistical results.

After determining which model was a best fit for each region, we looked at the factors within the winning models that significantly contributed to the fit (see *Methods*), and whether they were positive or negative, to understand the models’ relationship to the data (Figure 5a).

**Figure 5.**
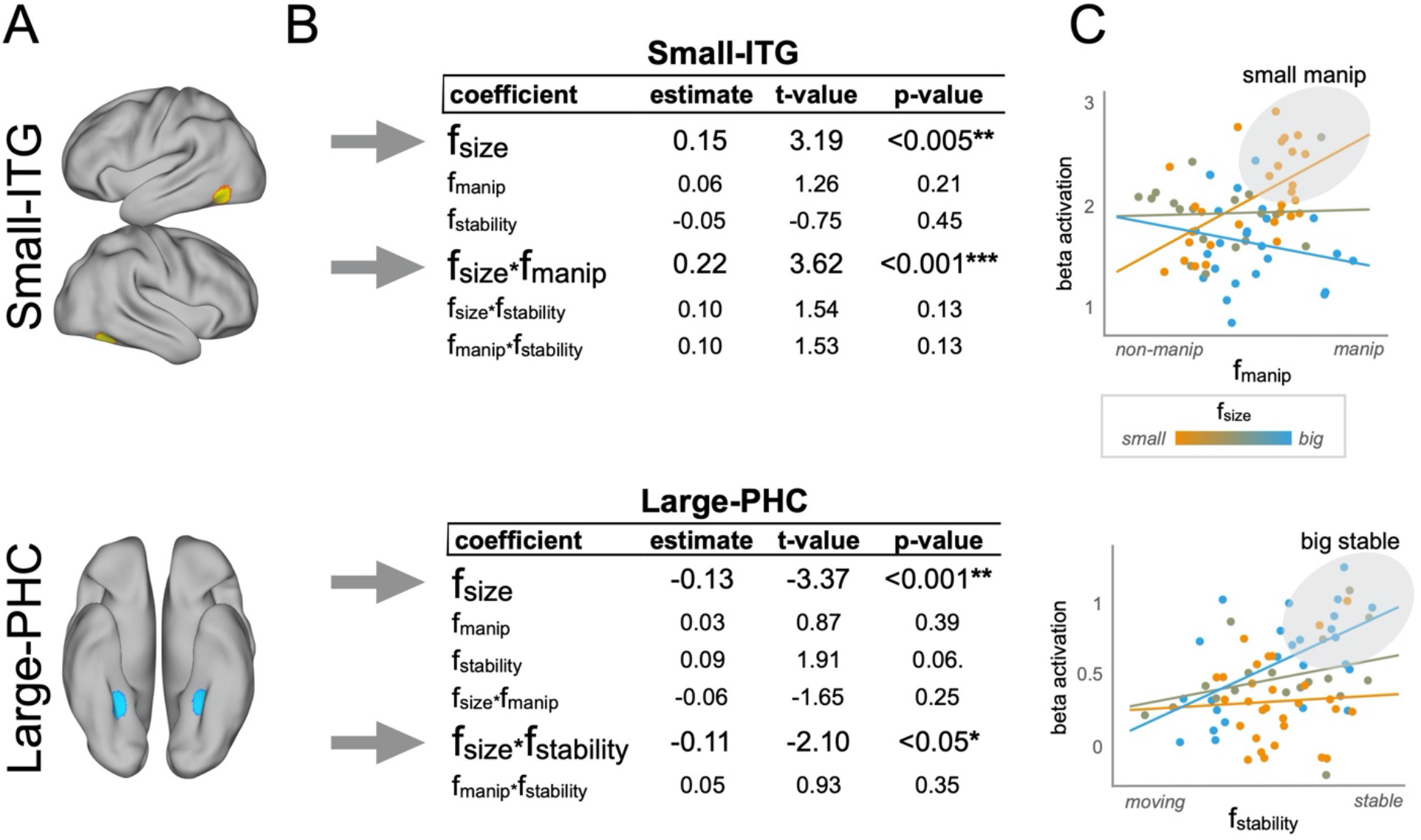
ROI analysis. a) Visualization of size ROIs Small-ITG and Large-PHC in one example subject. b) GLMs with ROIs’ activity as dependent variable and factor scores as independent variables. Shown are estimates for all predictors included in the model, t-value and p-value. The t-value was measured by dividing the coefficients by their standard errors. The p-value tested the hypothesis of obtaining the observed t-value if the coefficient were actually zero.c) Top: scatterplot of Small-ITG’s activation by f_manip_ (x-axis) and f_size_ (y-axis). Dots are color-coded based on the f_size_ weight from blue (low-weight, large size) to orange (high weight, small size). Bottom: scatterplot of Large-PHC’s activation by f_stability_ (x-axis) and f_size_ (y-axis). Same color-coding scheme as described for the plot above.

**Figure 6.**
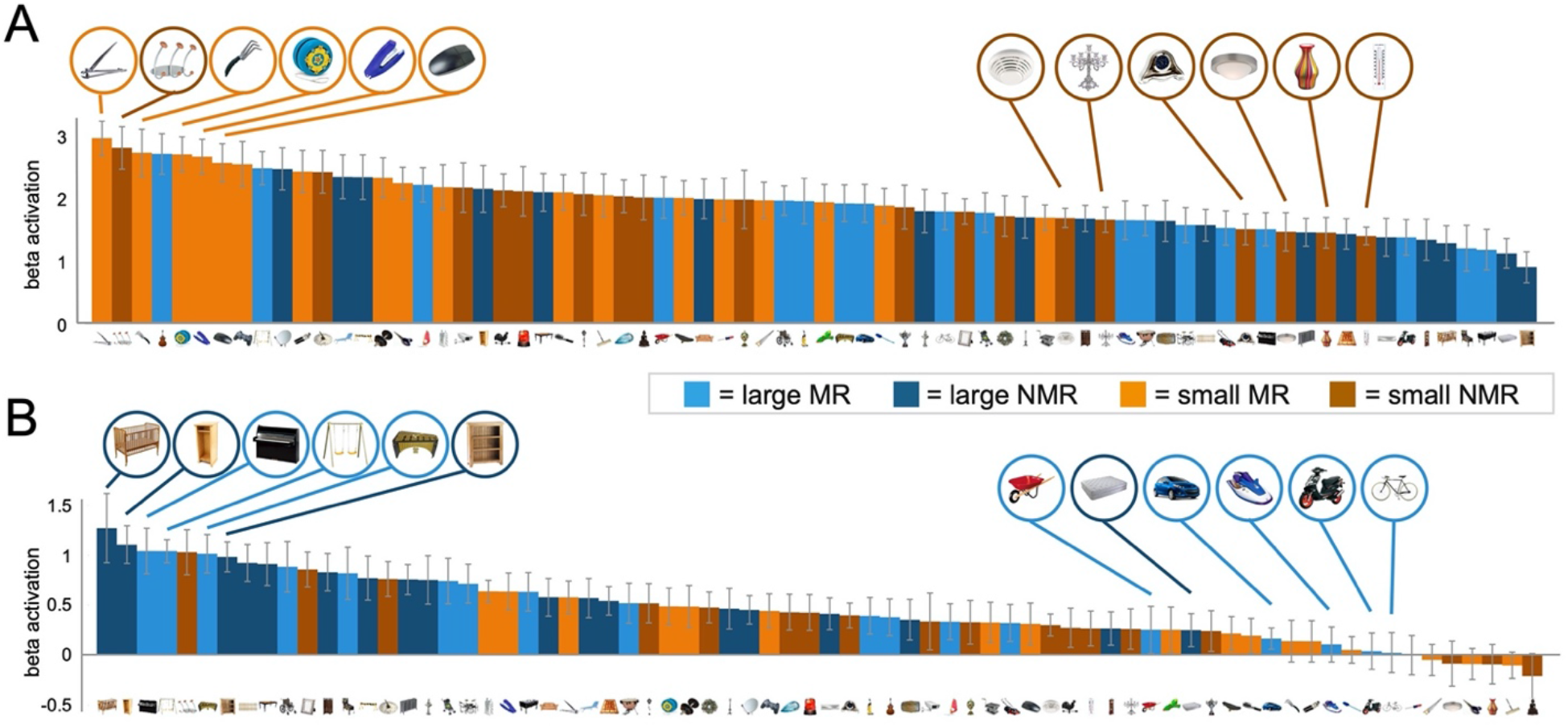
Barplots of ROIs activation. Mean activation to each of the 72 object categories are plotted for each region (a: Small-ITG; b: Large-PHC), where the categories are rank-ordered from most to least active. Object categories are color-coded based on their size and motor-relevance conditions (light orange: small motor-relevant; dark orange: small non-motor-relevant; light blue: large motor-relevant; dark blue: large non-motor-relevant). Error bars reflect ±1 SEM across subjects.

In Small-ITG, the results indicate that the best prediction of neural response magnitude was given by a combination of *size* and of the *manipulability* component of motor-relevance: there was a significant effect of f_size_, as well as the interaction of f_size_and f_manip_(see Figure 5a, top for statistical results). The interaction between these factors further indicates that it is specifically small objects involved in hand-object actions which drive the strongest responses in this region. Thus, the distinction of hand-object action and manipulability is a better descriptor of the responses in this region over the more general concept of motor-relevance.

In Large-PHC we observed a different pattern: the coefficients that significantly explained the model were f_size_ and the interaction of f_size_and f_stability_(see Figure 5b, bottom row for statistical results). Thus, responses in this region were driven most by large objects that are stable in the environment. This result in Large-PHC clarifies what we observed in earlier analyses employing the 2 x 2 design: rather than the low-action component of motor-relevance, it might be the *fixedness* component of this construct the one most driving neural representations in this area. That is, the *stability* component of non-motor-relevance is the one to best predict response magnitude in Large-PHC.

To better understand these relationships, we visualize them in Figure 5c: for each ROI, the activation to each category is plotted on the y-axis as a function of the manipulability factor for Small-ITG, and the stability factor for Large-PHC. Each dot is color-coded based on the f_size_ value for the corresponding object. This plot reveals that the manipulability factor drives activation in Small-ITG but mostly for small objects (increasing activity for more manipulable small objects, see Figure 5c, top). In contrast, inspection of the scatterplot for Large-PHC suggests that activation is driven by stability but mostly for large objects (increasing activity for more stable large objects, see Figure 5c, bottom). These relationships are also evident in the activation profiles of the two ROIs in Figure 6: For Small-ITG, tools are the small objects that drive the most activation (e.g. nail clipper, hand rack, yo-yo; see Figure 6a), while for Large-PHC, means of transport tend to be the large objects that drive the least activation (e.g. bike, scooter, jet-ski; see Figure 6b).

Taken together, these analyses begin to characterize the nature of the ecological covariation between size and motor-relevance, clarifying how these object properties fit together. For example, no one factor alone (size, manipulability, stability) best accounted for the responses in any ROI. Rather, manipulability (or hand-relevance) of an object matters more for small objects in the lateral cortex, and stability of an object matters more for large objects in ventral cortex. Finally, these findings are consistent with prior literature that has focused on either the ventral or lateral surface separately (e.g., Bracci et al., 2012; Mullaly and Maguire, 2011), unifying these separate observations in one common data set and analysis procedure.

#### 3.3.3 Additional ROIs

To more directly relate our findings to previous literature, we also conducted these same analyses in the classic scene regions, parahippocampal place area and occipital place area (PPA and OPA; Epstein et al., 2014), and object-responsive lateral occipitotemporal cortex (LOC; see Methods and Supplementary Material). These regions are adjacent and partially overlapping with the size ROIs (see also Konkle and Oliva, 2012). The findings in these classic regions generally converge with what was observed for the size ROIs: in LOC, there was a significant contribution of f_size_, and a significant contribution of the interaction between f_size_ and f_manip_. In PPA, there was a significant contribution of the size factor, and marginally of stability and of the interaction between size and stability (see Supplementary Material).

Further, we conducted these analyses in an additional occipitodorsal region identified with the [Large > Small] size contrast from two runs of the main experiment (Large-TOS) and in adjacent scene-selective occipital place area (OPA; see Supplementary Material). In Large-TOS, we found a result parallel to our observation in PPA: a significant contribution of f_size_ and of the interaction between f_size_ and f_stability_. However, this result did not replicate in OPA where only f_size_ had a significant contribution.

#### 3.3.4 Searchlight Analysis

By focusing on the ROIs produced by the size contrast, we were able to explore whether other factors beyond size and motor-relevance were important in these ROIs—to which the answer is clearly yes: motor-relevance was indeed split into two sub-dimensions, one more related to manipulability, and one more related to stability, which dissociated in explaining small- and large-object ROIs. However, by performing a targeted ROI analysis, we might have missed additional regions with different associations between these factors.

Given the complexity of the number of possible interactions among these factors, we approached the searchlight analysis with a more targeted comparison. Specifically, we mapped which regions showed a better model fit among two candidate models: one that associates size and manipulability (manip-model) and one that associates size and stability (stability-model; see Methods). All searchlight analyses were conducted within a reliable voxel mask (see Methods). The results of this analysis are shown in Figure 7.

**Figure 7.**
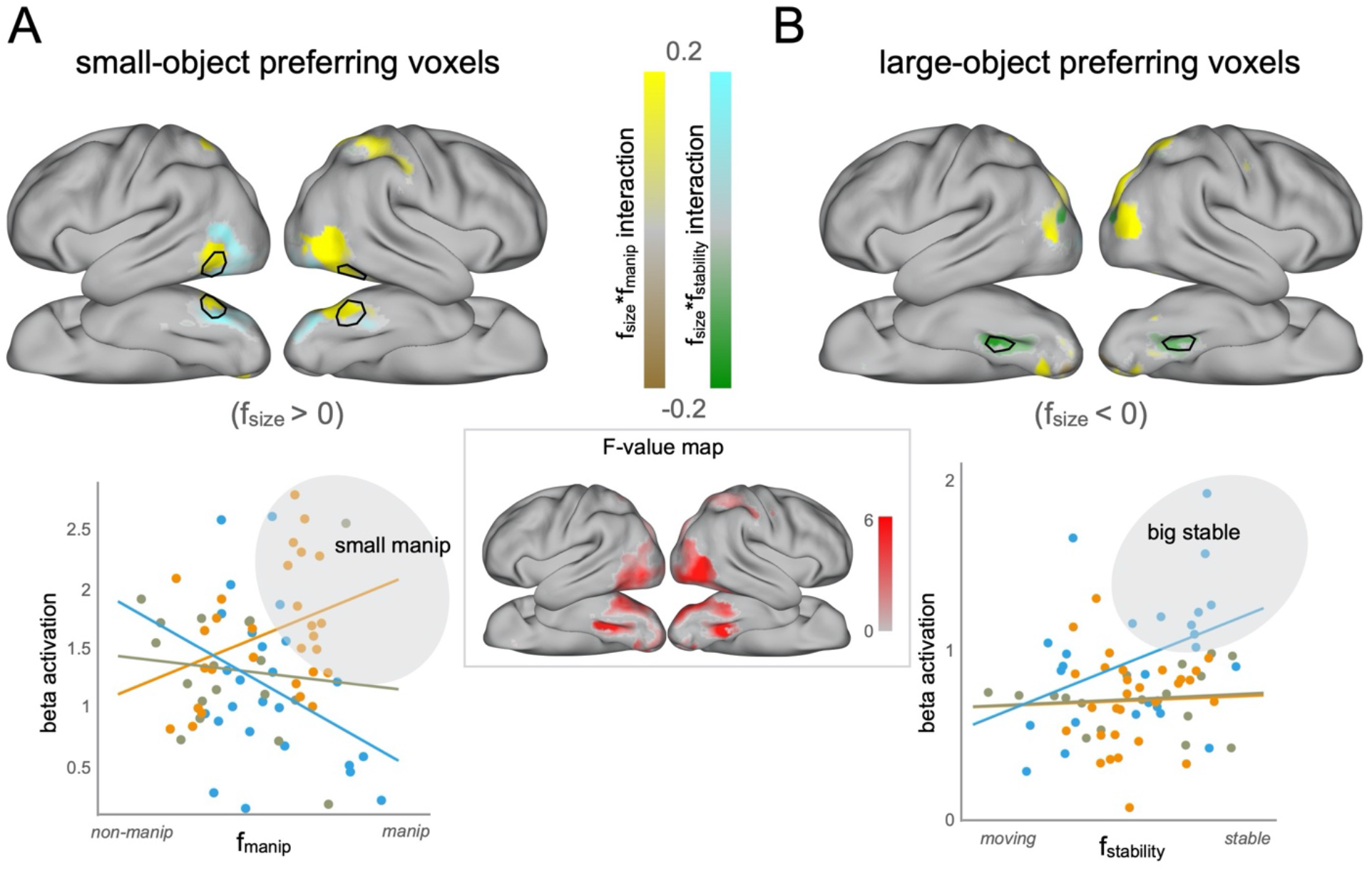
Searchlight Analysis. a) Top: surface map of interaction coefficients for small-object preferring voxels (i.e., voxels with a positive f_size_ coefficient). In voxels in which the winning model was the manip-model, that voxel is colored according to its f_size_*f_manip_ interaction coefficient estimate in a brown to yellow scale. In voxels in which the winning model was the stability-model, that voxel is colored according to its f_size_*f_stability_ interaction coefficient estimate in a green-to-cyan scale. Bottom: scatterplot of activation by f_manip_ (x-axis) and f_size_ (y-axis) for all voxels showing a positive f_size_*f_manip_ interaction in lateral OTC. Dots are color-coded based on the f_size_ weight from blue (low-weight, large size) to orange (high weight, small size). b) Top: surface map of interaction coefficients for large-object preferring voxels (i.e., voxels with a negative f_size_ coefficient). In voxels in which the winning model was the manip-model, that voxel is colored according to its f_size_*f_manip_ interaction coefficient term in a brown-to-yellow scale. In voxels in which the winning model was the stability-model, that voxel is colored according to its f_size_*f_stability_ interaction coefficient estimate in a green-to-cyan scale. Bottom: scatterplot of activation by f_stability_ (x-axis) and f_size_ (y-axis) for all voxels showing a negative f_size_*f_manip_ interaction in ventromedial OTC. Dots are color-coded based on the f_size_ weight from blue (low-weight, large size) to orange (high weight, small size).

From this analysis, we can gather two main conclusions. First, we found that the voxels with best fitting models (i.e., the ones with highest F-value, see Figure 7, center) focused mostly in the lateral occipitotemporal region and in the ventromedial occipitotemporal regions, in areas mostly overlapping with size ROIs Small-ITG and Large-PHC (outlined in black in Figure 7a,b). Thus, our targeted ROIs seemed to cover most OTC areas where voxels’ information could be explained by a combination of the three factors.

Second, we found that the best-fitting voxels resulting from the searchlight analysis largely confirmed our targeted ROI analyses. Indeed, small-object preferring voxels in the lateral occipitotemporal region (i.e., the ones with positive f_size_ coefficients for the winning model) presented a positive interaction of the size and manipulability factors (Figure 7a top, colored in yellow), similarly to what was observed in the ROI analysis for Small-ITG: that is, the smaller f_size_ (the smaller the object), and the higher f_manip_ (the more manipulable the object), the stronger the activation. Similarly, we found that large-object preferring voxels in ventromedial occipitotemporal region (i.e., the ones with negative f_size_ coefficients for the winning model) presented a negative interaction of the size and stability factors (Figure 7b, top, colored in green), similarly to what observed in our ROI analysis for Large-PHC: that is, the lower f_size_ (the bigger the object), and the higher f_stability_ (the more stable the object), the stronger the activation. For ease of comparison, we have drawn a black outline on the searchlight coefficient maps of the size ROIs Small-ITG (Figure 7a) and Large-PHC (Figure 7b) for one example subject, from which it is clear that the targeted ROI analyses do in fact highlight the major structure in neural responses by our three factors.

At lower F-value levels, we observed two more nuanced relationships. However, given the lower fit of either manip-model or stability-model within these voxels, one can only draw tentative inferences from these results.

First, a significant portion of small-object preferring voxels in the lateral occipitotemporal surface (Figure 7a, top, colored in cyan) are fit best by models with size and stability, and show a negative interaction between the size and stability factors. Inspection of activation to all 72 object categories in this area shows a preference for moving over non-moving things (see Supplementary material) which is in line with the presence of movement-preferring region MT+ in the lateral surface.

Second, we also observed large-object preferring voxels in the dorsal occipitoparietal region, where most voxels (Figure 7b, top, colored in yellow) indicate a positive interaction of the size and manipulability factors while a few voxels (colored in green) indicate a negative interaction of the size and stability factors. However, inspection of the scatter plots for these regions showed that these interactions were not as simple and easily interpretable as the ones observed in areas neighboring our size ROIs (see Supplementary Material). From these results, it is clear that there are systematically varying object responses in this dorsal occipital part of the cortex, but none of the three factors or their interactions cleanly provides a description of what explains the object activations.

## 4. DISCUSSION

Here, we examined whether the large-scale organization of neural responses to object size could be better explained by the factor of motor-relevance. This was not the case – we found that the contrasts of small vs large objects elicited a clear size organization for both motor-relevant and non-motor relevant objects. We also found that motor-relevance, independent of size, accounted for some of the large-scale structure of neural responses. This observation confirms the importance of motor interaction as an explanatory dimension in the ventral visual stream. Crucially, however, further analyses showed the two dimensions to not be independent, in that more typical combinations (small and motor-relevant, large and non-motor-relevant) elicited more reliable representations than less typical ones.

Our subsequent exploratory analyses helped to refine and clarify these results. Specifically, a factor analysis revealed that the construct of motor-relevance could be divided into two underlying dimensions, one better related to *manipulability* (i.e., related to hand performed actions), and one better related to an object’s *physical presence* and stability. Further ROI and searchlight exploratory analyses revealed that the manipulability factor was a better descriptor of activation in small-object preferring lateral OTC, while the stability factor was a better descriptor of activation in large-object preferring ventromedial OTC.

Broadly, these results begin to characterize how the ecological covariation between size and behaviorally relevant object properties contributes to the observed dissociations between object categories in the ventral stream regions with a preference for the inanimate domain. In the next two sections, we situate these findings in the literature, and discuss how they inform the deeper theoretical question of what drives the neural organization of inanimate objects.

### 4.1 Dissociations within the inanimate-domain

Objects are characterized by numerous visual properties, real-world size being one of them. However, size tends to correlate with other visual and non-visual properties. There are suites of partially related factors that predict neural responses to inanimate objects, with a major division between small, manipulable objects on one side and large, stable, navigationally relevant objects on the other (e.g., He et al., 2013; Peelen et al., 2013; Julian et al., 2016; Mullally and Maguire 2011; Troiani et al., 2014; Auger et al., 2013; Bracci et al., 2012; MacDonald and Culham, 2015; Bainbridge and Oliva, 2015). Overall, these findings point to a richer organization than simply object size as such, where different brain regions may be sensitive to size in the presence of other object properties for different reasons (Bi et al., 2015).

For example, along lateral OTC and middle temporal gyrus, there is a mosaic of partially overlapping regions whose response magnitudes are predicted by a range of related factors, from more primitive object shape (Grill-Spector, 2003), to real-world size (Konkle and Oliva, 2012; Julian et al., 2016), to tools and other objects that are manipulated by the hands (Lewis 2006; Bracci et al., 2012; MacDonald et al., 2015; Mruczek et al., 2013; Lingnau and Downing, 2015). These regions are also adjacent to motion-selective cortex and body-preferring regions (Grill-Spector and Weiner, 2014), including a region that jointly responds to both tools and hands, spanning the animate/inanimate divide (Bracci and Peelen, 2013; Striem-Amit et al., 2017). While the number of regions and exact functional divisions along this cortex are still being clarified, from our results and these others, it is clear that these regions are most strongly related to small, movable, manipulable items.

Along ventromedial OTC, where scene-preferring PPA is located, there are now a number of converging results characterizing the object properties that are best associated with responses to isolated objects. These factors include the degree to which the object invokes a local tridimensional space, its size, its tendency to stay fixed in the world, and can be generally summarized by an overarching construct of spatial stability and landmark suitability (Troiani et al., 2014, Mullally and Maguire, 2011; Julian et al., 2016; Auger et al., 2013; see also Epstein et al., 2014). While these studies examining PPA responses differ in their stimulus sets, designs, and even mode of presentation (e.g., pictures, mental imagery), they all leverage condition-rich designs and factor analyses to characterize the configuration of related object properties. While our observation of a role of spatial stability in PHC cannot in isolation lead to strong inferences, given that it is the product of a post-hoc analysis, it joins this collection of studies, adding to the diversity of stimulus sets and neuroimaging designs that show converging evidence that size in the context of position fixedness best explains the object responses in this region.

Our results, jointly with these collections of studies, suggest that a division based on manipulability for small objects and stability for large objects is fundamental in the organization of inanimate object responses.

### 4.2 What drives this object organization?

Turning to the broader question of *what drives* the spatial organization of object domains in visual cortex, as opposed to what visual properties are explicitly encoded in domain-specific regions, one answer is that the observed specialization is the result of evolutionary pressures to maximize the efficient mapping of visual representations onto the appropriate downstream regions engaged in action relevant processing for a given object domain (Caramazza and Shelton, 1998; Mahon et al., 2009; Mahon and Caramazza, 2011; Konkle and Caramazza, 2017; Conway, 2018). Candidate domains are manipulable (tools) and navigation-relevant objects. On this account, the representations computed in these regions are shape configurations and texture values that statistically reflect the distinguishing properties of manipulable versus navigation-relevant object domains. And, in the measure to which real-world size is differentially correlated with the two object domains (for example, large objects tend to be bulkier) it will have contributed to the evolving preferences for different visual features and configurations in the lateral and ventral occipitotemporal cortex. In other words, the two inanimate-object-preferring regions respond preferentially to, and presumably encode, the types of shape and texture properties that are associated with prototypical manipulable (small, graspable) objects and navigation-relevant (large, stable/stationary) objects.

It is relevant to note here that this view is fully compatible with accounts that emphasize the statistics of visual experience, wherein experienced eccentricity has played an evolutionary role in helping determine the localization of object domain preferences in the brain since it reflects the natural distribution of object viewing: more foveal, required for accurate reaching/grasping for small manipulable objects, versus more peripheral, associated with the requirement for capturing spatial (context) relations for large space-defining objects (Malach et al., 2002, Arcaro et al. 2009; Mahon and Caramazza, 2011; Konkle and Oliva, 2012; Gomez et al., 2017).

A crucial distinction is being drawn here between, on the one hand, the factors that may have determined the observed neural specialization in different regions of OTC for different types of inanimate objects and, on the other hand, the information being computed/represented in those specialized areas. The role of the object property “size” at these two levels of description of neural specialization is illustrative in this regard. It could be argued that real-world object size, because of its role in distinguishing between manipulable and navigation-relevant object types, contributed to help select the visual shape/texture properties that characterize the observed domain specialization in OTC. But what about the role of the visual property “size” in characterizing the computations/representations in these neural regions? Is this visual property explicitly computed/represented in these areas? One reading of the available results is that size is not directly computed/represented in these areas.

There are at least three senses of the property size in the context of visual processing: subtended visual angle, perceived physical size, and real-world size. Konkle and Oliva (2012) found that only objects’ real-world size, and not retinal or inferred size, is related to regional specialization. However, this generalization is tempered by the observation, reported in Konkle and Caramazza (2013), that the effect of real-world size is limited to inanimate objects: real-world size differences in the animate domain are not associated with regional specialization. An implication of this observation, consistent with the results reported here, is that it is not real-word size, as such, that drives neural responses but the types of visual shapes that are typically associated with inanimate small, manipulable objects versus large, stable objects (e.g., Long et al., 2016). Furthermore, the property real-world size of an object is a “constructed” value that is retrieved from memory and has no obvious description in a vocabulary of visual properties such a shape, orientation, texture, and color. Indeed, the notion “real-word size” is a kind of conceptual/functional information like the properties “manipulability” and “navigation-relevance”, and none of these types of information is directly represented in the putative, size-responsive areas.

By positing a key role for domain-related processing, this view provides an account of data from congenitally blind individuals, who show response preferences in similarly located regions of occipitotemporal cortex when hearing the names of animate versus inanimate objects or big versus small artifacts (e.g., Mahon et al., 2009; Wolbers et al., 2011; He et al., 2013; Peelen et al., 2013; Bi et al., 2015; Mattioni et al., 2020). Since ontogenetic visual experience could not have contributed to the similar patterns of object-preferring effects in high-level visual areas in congenitally blind and sighted individuals, it invites the conclusion that the observed specialization predates such experience. While the domain account provides a principled framework for explaining the potential functional basis for the observed large-scale organization of “visual” cortex, it is silent on the specific representational content at any given level of visual processing. Investigation of the specific perceptual features – e.g., elongated versus bulky shapes, texture properties, curvature patterns, or specific combinations of such features – that are represented/computed in the various object-type-preferring regions is an active, promising area of research (e.g., Andrews et al., 2010; Rajimehr et al., 2011; Bracci and Op de Beeck, 2016; Long, Yu and Konkle, 2018), aided by modeling with deep CNNs (e.g., Güçlü and Van Gerven, 2015; Khaligh-Razavi and Kriegeskorte, 2014; Yamins et al., 2014; Kubilius et al., 2016).

## Acknowledgments

This work was supported in part by a grant from the Fondazione Cassa di Risparmio di Trento e Rovereto to the CIMeC – Centro Interdipartimentale Mente/Cervello, University of Trento (AC) and by the National Institutes of Health, National Eye Institute (Fellowship F32EY022863-01A1 to TK) and was conducted at the Laboratory for Functional Neuroimaging Center at the Center for Mind/Brain Sciences (CIMeC), University of Trento.

