## Supplemental Material for "The contribution of object size, manipulability, and stability on neural responses to inanimate objects"

### Supplementary Material

#### Supplementary ROIs Analyses
#### Methods

***Large-TOS and Categorical ROIs selection.*** In addition to the small-ITG and large-PHC, we identified left and right large-object selective TOS (Large-TOS, see Figure S1a; n=15 for left hemisphere, n=13 for right hemisphere; average size of 226 voxels).

To connect our main findings with previous literature, we also defined gray-matter masked spherical ROIs (12mm radius) around the peak of activation for object-selective LOC and scene-selective PPA and OPA. PPA and OPA were localized in each participant using the contrast of Scenes > Objects from the functional Localizer runs (n=18), LOC was localized in each participant using a contrast of Objects > Scrambled from the same runs (n=17). The average size was 228 voxels for LOC, 257 voxels for PPA and 248 voxels for OPA. LOC was localized in all but one subject due to a technical issue related to stimuli presentation.

***ROIs overlap.*** When comparing the location of the spherical ROIs, there was systematic overlap between Small-ITG and LOC, between Large-PHC and PPA and between Large-TOS and OPA (see also Konkle & Oliva, 2012). To quantify this, ROI overlap was computed by extracting, for each subject, the number of voxels shared between Small-ITG and LOC, between Large-PHC and PPA and between Large-TOS and OPA. The average of the shared voxels was then computed across subjects and divided by the average extension of LOC, PPA or OPA across subjects. On average, 33% of left LOC overlapped with left Small-ITG and 8% of right LOC overlapped with right Small-ITG: right Small-ITG was located more anteriorly and dorsally than its left counterpart, while left and right LOC were relatively symmetrical. Additionally, 47% of left PPA overlapped with Large-PHC while 50% of right PPA overlapped with right Large-PHC. Finally, 13% of left OPA overlapped with left Large-TOS and 22% of right OPA overlapped with right Large-TOS.

**Results**

***LOC and PPA.*** We observed a pattern of results mostly convergent with our main results when running regression models in LOC and PPA $f_{size}$, $f_{manip}$ and $f_{stability}$ as predictors (see Figure S1 and S2).

Similar to what observed with Small-ITG, LOC was significantly associated to $f_{size}$, $f_{manip}$ and their interaction (see Figure S1). This result differs only slightly from small-ITG, in which there was no significant contribution of $f_{manip}$ alone.

PPA was significantly associated to $f_{size}$ and marginally to the interaction between $f_{size}$ and $f_{stability}$ (see Figure S2). This result is largely in line with the one observed in Large-PHC.

**Figure S1.**

**LOC**

| coefficient | estimate | t-value | p-value |
|---|---|---|---|
| $f_{size}$ | 0.14 | 2.8 | <0.01** |
| $f_{manip}$ | 0.12 | 2.5 | <0.05* |
| $f_{stability}$ | -0.02 | -0.33 | 0.74 |
| $f_{size}$*$f_{manip}$ | 0.27 | 4.10 | <0.001*** |
| $f_{size}$*$f_{stability}$ | 0.13 | 1.83 | 0.07 |
| $f_{manip}$*$f_{stability}$ | 0.13 | 1.84 | 0.07 |

**Figure S2.**

**PPA**

| coefficient | estimate | t-value | p-value |
|---|---|---|---|
| $f_{size}$ | -0.11 | -3.28 | 0.005** |
| $f_{manip}$ | 0.01 | 0.37 | 0.71 |
| $f_{stability}$ | 0.09 | 1.90 | 0.06. |
| $f_{size}$*$f_{manip}$ | -0.05 | -1.2 | 0.23 |
| $f_{size}$*$f_{stability}$ | -0.09 | -1.90 | 0.07. |
| $f_{manip}$*$f_{stability}$ | 0.01 | 0.25 | 0.80 |

***Large-TOS and OPA.*** We found a significant contribution of $f_{size}$ and of the interaction between $f_{size}$ and $f_{stability}$ in Large-TOS (Figure S3b), but inspection of scatterplot and barplot of activation to individual objects didn't reveal a clear direction of the interaction (Figure S3c and Figure S4). Finally, OPA was only significantly associated to $f_{size}$, without other factors or interactions significantly contributing (Figure S5).

**Figure S3.**

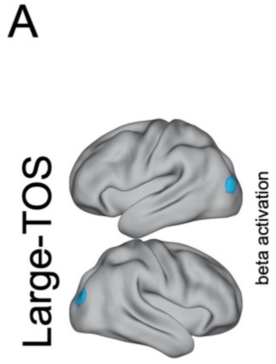


B

**Large-TOS**

| coefficient | estimate | t-value | p-value |
|---|---|---|---|
| $f_{size}$ | -0.12 | -2.46 | <0.05* |
| $f_{manip}$ | 0.05 | 1.06 | 0.29 |
| $f_{stability}$ | -0.06 | -0.93 | 0.35 |
| $f_{size}*f_{manip}$ | 0.005 | 0.08 | 0.94 |
| $f_{size}*f_{stability}$ | -0.16 | -2.20 | <0.05* |
| $f_{manip}*f_{stability}$ | 0.02 | 0.23 | 0.82 |

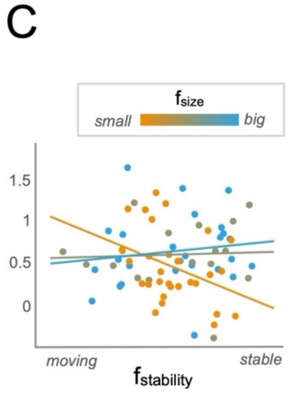


**Figure S4.**

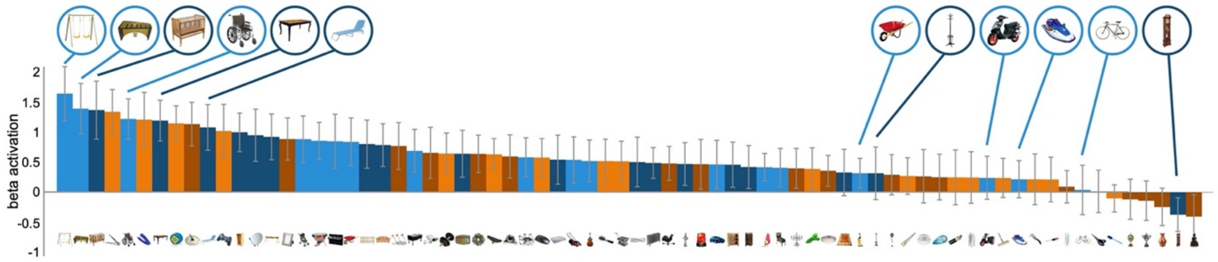


**Figure S5.**

**OPA**

| coefficient | estimate | t-value | p-value |
|---|---|---|---|
| $f_{size}$ | -0.11 | -2.17 | <0.05* |
| $f_{manip}$ | 0.04 | 0.87 | 0.39 |
| $f_{stability}$ | 0.03 | 0.50 | 0.61 |
| $f_{size}*f_{manip}$ | 0.00 | 0.03 | 0.98 |
| $f_{size}*f_{stability}$ | -0.07 | -0.96 | 0.34 |
| $f_{manip}*f_{stability}$ | 0.08 | 1.15 | 0.25 |

**Supplementary Searchlight Analyses**
When performing the searchlight analysis, we observed additional sub-regions which we did not cover in the main analyses, as the model fits tended to be lower. Here, we plot the activation within these sub-regions for each of our 72 objects.

Figure S6a depicts the sub-region composed of small-object preferring voxels which show a positive $f_{size}$*$f_{stability}$ interaction, showing a preference for moving over stable big objects, and an overall trend in preferring moving over non-moving across object size. This area might be partly overlapping with region MT+ which is known to show a preference for moving over non-moving stimuli.

Figure S6b depicts the sub-region composed of large-object preferring voxels which show a negative $f_{size}$*$f_{stability}$ interaction, showing a preference for stable over moving big objects. Curiously, this region seems to be partly overlapping with Large-TOS and OPA.

Figure S6c depicts the sub-region composed of large-object preferring voxels which show a positive $f_{size}$*$f_{manip}$ interaction, showing a preference for manipulable over non-manipulable small objects, and a trend towards preferring non-manipulable over manipulable small objects.

**Figure S6**

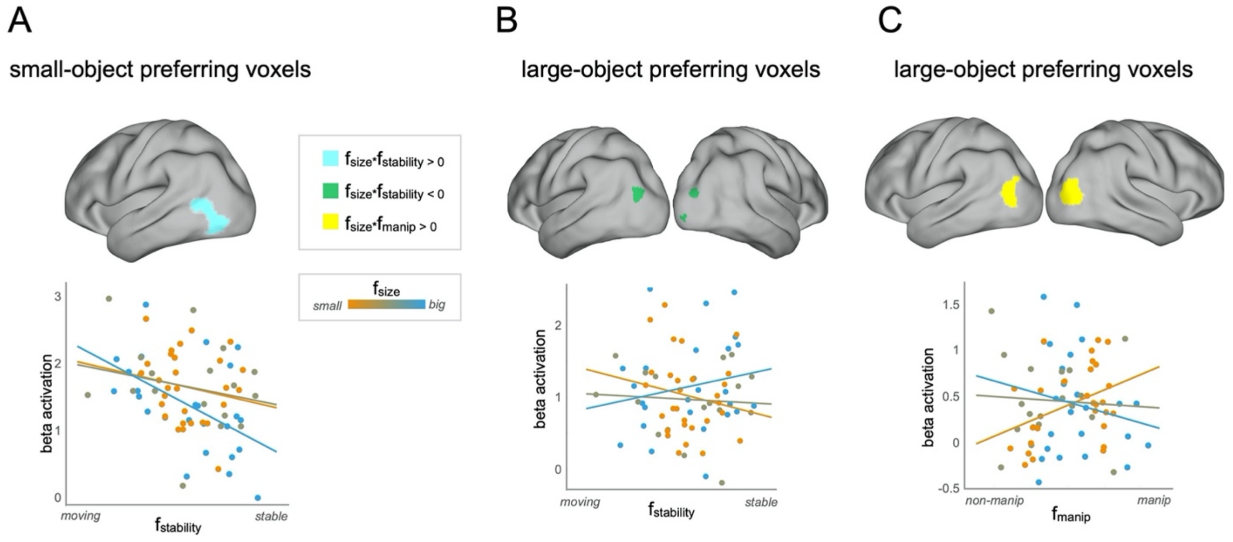
